# In Silico Mechanobiochemical Modeling of Morphogenesis in Cell Monolayers

**DOI:** 10.1101/189175

**Authors:** Bahador Marzban, Xiao Ma, Xiaoliang Qing, Hongyan Yuan

## Abstract

Cell morphogenesis is a fundamental process involved in tissue formation. One of the challenges in the fabrication of living tissues in vitro is to recapitulate the complex morphologies of individual cells. Despite tremendous progress in understanding biophysical principles underlying tissue/organ morphogenesis at the organ level, little work has been done to understand morphogenesis at the cellular and microtissue level. In this work, we developed a 2D computational model for studying cell morphogenesis in monolayer tissues. The model is mainly composed of four modules: mechanics of cytoskeleton, cell motility, cell-substrate interaction, and cell-cell interaction. The model integrates the biochemical and mechanical activities within individual cells spatiotemporally. Finite element method (FEM) is used to model the irregular shapes of cells and to solve the resulting system of reaction-diffusion-stress equations. Automated mesh generation is used to handle the element distortion in FEM due to the large shape changes of the cells. The computer program can simulate tens to hundreds of cells interacting with each other and with the elastic substrate on desktop workstations efficiently. The simulations demonstrated that our computational model can be used to study cell polarization, single cell migration, durotaxis, and morphogenesis in cell monolayers.

## Introduction

Recent advances in tissue engineering such as 3D bioprinting technologies ^1^ have made it possible to create on demand 3D complex functional human tissues/organs. The ability to engineer 3D complex tissues/organs on demand will have unprecedented impact in regenerative medicine ^1^, disease modeling and drug discovery ^2^. In tissue engineering, especially for 3D bioprinting, the central challenge is to reproduce the complex micro-architecture of cells and extracellular matrices (ECM) in microscale resolution to recapitulate biological functions ^3^. A thorough understanding of how microtissue morphogenesis occurs in an engineered microenvironment is critical for the 3D bioprinting approach to succeed.

Tissue/organ morphogenesis is a complex process occurring at multiple scales. Focusing on the whole organ scale, considerable research has been devoted to elucidation of the physical principles underlying the formation of the overall morphologies of organs ^4–6^, as well as the nutrient consumption and transport in bioreactors ^7^ for tissue engineering. In these whole-organ level studies, information at the individual cell level has been homogenized or ignored. At the other extreme of the length scale, the genetic and molecular causes that dictate the tissue/organ formation have been intensively studied ^8^. There is gap between our understanding of how phenotypic morphologies at the organ level emerge from genetic information. Studies at the cellular and microtissue level play an indispensable role to bridge these two scales. Despite its importance, very little work has been done on the cell and microtissue morphogenesis.

The phenotypic morphologies of cells including cell shape and cytoskeleton architecture, cell-ECM and cell-cell adhesions, can be best seen by comparing four types of tissues: muscle tissue, nerve tissue, epithelial tissue, and connective tissue. Each of these different tissues exhibit characteristic morphologies in cell shape and cytoskeleton architecture. These four basic types of tissues are arranged spatially in various patterns (e.g., sheets, tubes, layers, bundles) to form organs. Gene expression only dictates what proteins to make and subsequently what biochemical reactions to carry out, the emergence of spatial morphologies must be determined by biomechanical principles and the coupling between biomechanics and biochemistry ^9–11^. Mechanobiochemical coupling is exemplified by the recent discoveries in the field of mechanobiology. Cellular functions including cell migration and cytoskeletal dynamics that are closely related to cell morphogenesis, have been shown to be regulated by various mechanical cues such as matrix elasticity ^12^, matrix topology ^13–18^, matrix dimensionality ^19–23^, cell-ECM/cell-cell adhesions ^24^, and cell shape constraints ^25–28^. Therefore, mechanical and geometric properties of cells and their microenvironments at the length scale comparable to single cells can have a dominant effect on the microscopic tissue morphology.

Mathematical models based on reaction-diffusion equations at the cellular scale were developed to understand spatial pattern formation in the context of cell migration, such as cell polarization ^29,30^ and cell morphogenesis ^31,32^. However, biochemical models lack the consideration of mechanotransduction thus cannot adequately capture cell morphogenesis. Biomechanics models were developed to interpret specific aspects of cell spreading, for example, the distribution patterns of traction force ^33^, cell adhesion ^34^, and cytoskeleton dynamics ^35–39^. sIn contrast, biomechanics models lack consideration of biochemical signaling and thus fail to account for biochemical regulations. A thorough understanding of cell and microtissue morphogenesis will require the elucidation of how the mechanical and biochemical events are spatiotemporally integrated at the cellular scale.

In this work, we first developed a 2D single-cell computational model. The single-cell model mainly integrates three modules: cell migration, cytoskeleton mechanics, cell-substrate interaction. We then extended the single-cell model to multicellular monolayer model by adding a module of the cell-cell interaction. Finite element method is used to solve the resulting system of partial differential equations and the model was implemented in an in-house MATLAB code.

## Model Description

### Physical and mathematical Domains

Four physical domains are defined for the cell: solid phase cytoskeleton domain, fluid phase cytosol domain, membrane domain, and nucleus domain. The elastic substrate (underneath the cell) domain is denoted by. In the cell membrane domain and cytosol domain, reaction-diffusion equations of diffusive molecules will be formulated to model the protrusion and retraction signals. In the cytoskeleton domain, solid mechanics equations will be formulated and the mechanical stresses experienced by the cell will be solved. The present computational model is concerned with the cell monolayer adhering to a flat substrate. Each cell is modeled as a two-dimensional (2D) continuum, which reflects the flatness of the lamellipodia for cells cultured on 2D flat substrates. Because 2D model of the cell is adopted, the physical domains , , , and can be described by the same mathematical domain, denoted by.

### Cytoskeleton module

#### Cytoskeleton mechanics

A rather simple mechanics model of cytoskeleton is adopted here. The motions of biological cells are at the low Reynolds number, inertia force can be neglected ^40^, so at each time instant, the cell can be considered in a quasi-static equilibrium. The equilibrium equation of the cytoskeleton in the domain Ω_CSK_ are written by using Cartesian tensor notation (summation over repeated indices is adopted hereinafter) as

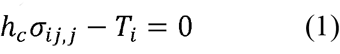

where *σ_ij_* is the Cauchy stress tensor of the cytoskeleton (*i* and *j* takes values of 1 and 2 in 2D), *h_c_* is the thickness of the cell and is assumed to be volume of the cell divided by the area of the cell. Volume conservation for the cell adopted in this model., *T_i_* is the traction stress exerted on the substrate by the cell. Traction stress is assumed to be linearly proportional to the displacement of the cell with respect to the substrate

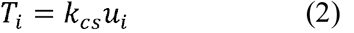

where *u_i_* is the displacement, *k_cs_* is the spring constant of the cell-substrate linkage that will be defined later in Eq. (18). At the cell edge (denoted by Γ) where there is no cell-cell adhesion, the stress-free boundary condition holds: *σ_ij_n_j_* = 0, where *n_j_* is the normal direction at the cell edge. In the present model, the cytoskeleton is composed of passive and active networks. For the sake of simplicity, here a simple elastic constitutive relation is adopted,

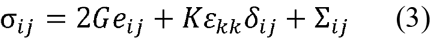

where *ε_j_* is the strain tensor, *e_ij_* = *ε_ij_* – *ε_kk_*/3 is the deviatoric strain tensor, *G* and *K* are shear and bulk modulus of the passive network, ∑_ij_ is the active isometric tensile stress (ITS) tensor from the active part of the cytoskeleton, which will be defined later in Eq. (5). Use of the small-strain Hooke’s law in Eq. (3) for the large deformation that occurs during cell motility deserves some explanations here. In this model, when solving the elasticity problem for a migrating cell, at each time instant we treat the current configuration as the stress-free state. This is an ad hoc treatment, but it can be understood as the dynamic bonds that forms the passive network of the cytoskeleton remodels fast enough to release the passive stress in the cytoskeleton.

#### Stress-fiber structure tensor

To account for the anisotropic fiber formation, a second-order tensor *S_ij_*, referred to as the stress-fiber structure tensor, is defined and its time evolution is described by

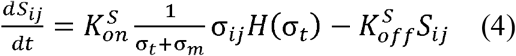

where *σ_m_*, 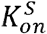 and 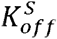 are model parameters, *σ_t_* = *σ_kk_*/2 is the cytoskeletal tension, *H*( ) denotes Heaviside function and is defined as: *H*(*x*)=1 when *x* > 0 and *H*(*x*)=0 when *x* ≤ 0. Denoting the maximal eigenvalue and the corresponding eigenvector of *S_ij_* by *λ*_1_ and 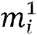, respectively, the ITS tensor ∑_*ij*_ is defined as

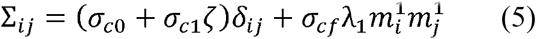

where *σ*_*c*0_, σ_*c*1_, and *σ_cf_* denotes the baseline, retraction signal-associated, and stress fiber-associated contractility, *ζ* is the concentration of retraction signal that will be introduced below.

The dyadic product of unit vector 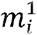 produces the tensor 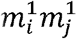, which has its only non-zero-principle-value principle direction along 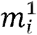

#### Cell motility module

Cell migration plays a pivotal role in tissue morphogenesis which is a fundamental process of the embryonic development ^41–43^. For in vitro 2D cell migration, the biomechanical machineries that drive migration, and the biochemical signaling networks that regulate the migration machineries have been studied intensively ^44^. The current understanding of cell migration is a synthesis from studies of different cell types ^45^. Single cell migration on 2D surfaces can be described as a coordinated and integrated process of different modules: cell polarization (i.e., front-and-rear formation), protrusion of lamellipodia/filopodia/lobopodia, formation of new adhesions in the front, releasing of aging adhesions at the rear, and cytoskeleton or membrane skeleton contraction to move the rear forward ^41,44^.

The intracellular biochemical signaling have been interpreted as to form networks and constitute an internal excitable system ^46,47^ Cells with this internal excitable system are able to spontaneously polarize and make persistent random walks in the absence of external guidance cues ^31^, and to carry out directed movement when biased by external signal gradients (i.e., chemotaxis). In recent years, it has been shown that cell migration can also be guided by a variety of mechanical cues of the microenvironment, such as substrate rigidity (durotaxis ^12^), shear flow (mechanotaxis ^48^), interstitial flow (rheotaxis ^49^), and cell-cell contact force (plithotaxis ^50^).

In this work, we adopt a similar modeling concept as Satulovsky et al. ^31^ where a few phenomenological variables are used to represent the concentration of various proteins involved in cell migration. The cell migration model is described below.

#### Reaction-diffusion of protrusion and retraction signals

Previous studies indicated that the active forms of protrusion and retraction signals are membrane-bound proteins ^32,51^. Two phenomenological variables *ξ* and *ζ* are defined in the physical domain of the membrane Ω_Mem_ to account for the concentration of active form of protrusion (e.g., Rac, Cdc42) and retraction (e.g., ROCK) signals, respectively ^31^. Here variables *ξ* and *ζ* are scaled by their saturation values respectively and are in units of μm^−2^. This means the maximal value *ξ* and *ζ* can reach is 1 µm^-2^. Their time evolution equations are defined as

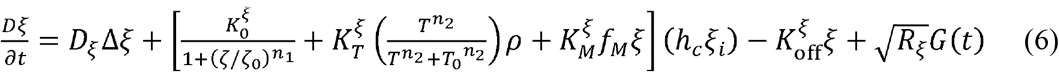

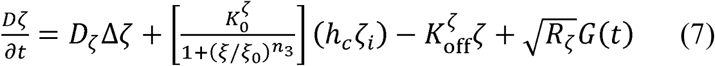

where 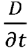 represents the material time derivative (the cytosol is assumed to be moved with the cytoskeleton, no fluid dynamics is considered here),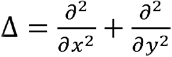 is the Laplace operator, *D_ξ_* and*D_*ζ*_* are the protrusion and retraction diffusion constants in the membrane, 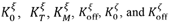 are rate constants, *ξ_0_*, *ζ_0_*, *n*^1^ *n*^2^, *n*_3_ are model parameters describing the autoinhibition relation between the protrusion and retraction signals, *R_ξ_* and *R_ζ_* denote strength of random noise for protrusion and retraction signals, respectively, *G*(*t*) is a Gaussian random process of mean zero and variance unity and 〈*G*(*t*)*G*(*t*’)〉 = *δ*(*t* — *t*’), *ξ*_*i*_ and *ζ_i_* are the concentration of the inactive forms of protrusion and retraction signaling molecules in the cytosol, which are in units of µm^-3^. The diffusion of inactive protrusion and retraction signaling molecules in the cytosol are considered much faster than in the membrane. To be simple, *ξ_i_* and *ζ_i_* are assumed to be uniform in the cytosol and are calculated by the following two equations, respectively,

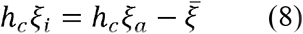

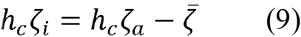

where *ξ_a_* and *ζ_a_* are model parameters denoting the total concentrations of both active and inactive forms for protrusion and retraction signals, respectively, 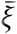 and 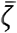 are the spatial mean values of *ξ* and *ζ*, respectively. In Eq. (8), multiplying *ξ_i_* by *h_c_* to convert a volume concentration to an area concentration is based on the assumption that the cell is flat and the diffusion in the cell thickness direction is instantaneous. Note that we update the cell thickness, *h_c_*, in each time step.

#### Movement of the cell (i.e., protrusion and retraction)

The movement of the cell domain is composed of two parts. One is the cell protrusion caused by the actin polymerization at the leading edge of the lamellipodia. The other is the passive retraction as a result of active cytoskeleton contraction. A protrusion velocity ***ν**^p^* is defined as a function of the protrusion signal at the cell edge *Γ* as

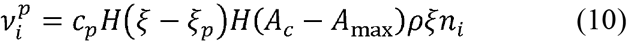

where *c_p_*, *ξ_p_*, and *A*_max_ are model parameters, *n_i_* is the outward normal unit vector at the cell edge, *A_c_* is the cell area, *H*( ) denotes Heaviside function. The retraction velocity is assumed to be proportional to the displacement *u_i_* of the cytoskeleton as,

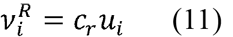

where *c_r_* is a model parameter. Note that the retraction velocity is applied to the whole cytoskeleton.

#### Cytoskeletal asymmetry

Experimental studies have implied that the cell polarity (i.e., head-and-tail pattern) is maintained through the long-lived cytoskeletal asymmetries including microtubules ^52^. To incorporate the cytoskeletal asymmetries in the model, a vector ***M*** is defined to represent the polarity of the asymmetric cytoskeleton and its time evolution equation is defined as,

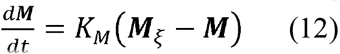

where *K_M_* is a model parameter, vector *M_ξ_* is a vector defined based on the protrusion signal,

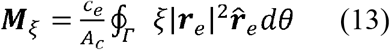

where 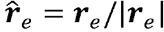 is a unit vector and ***r**_e_* is the position vector of the edge points relative to the center of the nucleus, |***r**_e_*| is the length of ***r**_e_*, *dθ* is the differential angle corresponding to the differential arc length, where *θ* is the angle coordinate of the edge point in the polar coordinate with the nucleus center as the origin. As implied by Eq. (12), in the steady-state (*d**M***/*dt*=0), the cytoskeleton asymmetry vector ***M*** is equal to the vector ***M**_ξ_*. The cytoskeleton-asymmetry function *f_M_*(*s*) in Eq. (6) is defined with the angle *θ_M_* of the vector ***M*** as,

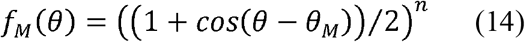

#### Nucleus deformation and movement

The nucleus is modelled as an elastic structure that deforms upon the compression of the cell membrane and moves with the cytoskeleton. In the present model, there are no mechanotransduction associated with the nucleus. Rather, the nucleus is a passive material and can resist deformation and contribute to the shape of the cell in cases cell are elongated or compressed. In the finite element-based numerical implementation, the nucleus is discretized into networks of springs. The velocity ***V*** of each node is calculated using the sum of the 3 forces, ***F***_ela_, ***F***_mem_, and ***F***_cyto_, divided by *μ*_nu_. ***F***_ela_ is the elastic recoiling force derived from the elastic deformation energy of the nucleus, meaning that both strain energy of the truss elements and the area energy of the nucleus, ***F***_mem_ is the compressive force exerted by the cell membrane when the membrane is within a cutoff distance from the edge of the nucleus, *F*_cyto_ is the friction force between the cytoskeleton and the nucleus at their interface.*μ*_nu_ is a viscosity parameter. the nodal velocity ***V*** is then used to update the position and the shape of the nucleus in the numerical integration procedure.

#### Cell-substrate interaction module

To incorporate the dynamic remodeling of focal adhesion, a phenomenological variable *ρ* is defined to describe the density distribution of focal adhesion-associated proteins (e.g., integrins, talins, vinculins, etc.), ranging from zero (no integrin-mediated cell-substrate adhesion) to one (mature FAs). The time evolution of *ρ;* is described by

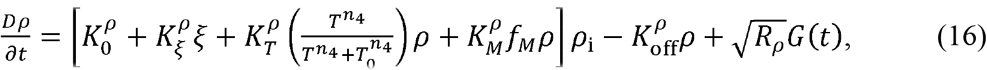

where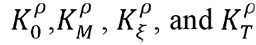 are the rate constants for the spontaneous, auto-activation, protrusion signal-dependent, and stress-mediated focal adhesion formation, respectively, 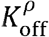 is a decay constant, *T* denotes the magnitude of the traction stress, *T*_0_ and *n*_4_ are model parameters, and *ρ_a_* represents the average density of the total amount of bound and unbound focal adhesion proteins, *R_ρ_* is the strength of random noise. Here the redistribution (e.g., via active transportation and passive diffusion) of unbound focal adhesion proteins is assumed to be faster than other time scale of focal adhesion formation, the unbound focal adhesion protein density *ρ_i_* in the cytosol is simply computed as.

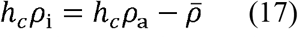

where 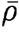 is the mean value of *ρ* in the membrane domain. Denoting *k_FA_* and *k_ECM_* the equivalent spring constants of the focal adhesion and the substrate, respectively, the spring constant of the cytoskeleton-substrate linkage is given as

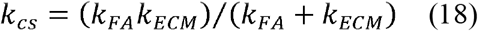

The mechanics of the cell is coupled to the dynamics of focal adhesion remodeling through the spring stiffness *k_FA_* by the following relation

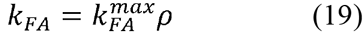

where 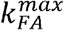 is the maximal stiffness when the focal adhesion density *ρ* = 1 μm^-2^ (i.e., mature focal adhesion).

#### Cell-cell interaction module

In cell monolayers where cells are connected mechanically by cell-cell adhesion, the static equilibrium of the cells depends on the cell-cell contact ^53–56^ in addition to cell-substrate adhesions. To simulate the dynamic process of formation and dissociation of cell-cell adhesion, a stochastic model is used to determine the binary state of the cell-cell adhesion as follows. When cell-A and cell-B is in close contact, the state of the cell-cell adhesion can be either “on” or “off”. The “on” state indicates that the cell-cell adhesion is established. The “off’ state indicates that although two cells are in close contact but they do not adhere to each other. The probability of the “off’ state per unit edge length and unit time is denoted by *φ*. when the cell-cell adhesion state is “on”, the stress between cell-A and cell-B, *P_i_*, is calculated as

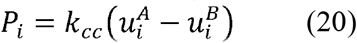

where 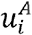 and 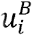 are the displacement of cell-A and cell-B at their edges (where the cell-cell adhesion is formed), respectively, and *k_cc_* is the spring constant of the cell-cell linkage.

#### Mechanobiochemical coupling

These different modules are coupled through the mechanics of the tissue. As illustrated in Fig. 1B, through molecular scale mechanotransduction pathways, mechanical stresses in the cell are converted into biochemical activities, which in turn regulate the assembly/disassembly of macromolecular of the cell. The macromolecular assembly and disassembly alter the structural, geometrical, and material properties of the cell, which, according to the continuum/structural mechanics theory, will subsequently change both the internal stress (cytoskeleton stress) and stress at the boundary (cell-matrix and cell-cell adhesion stress). Thus, mechanics of the cell, biochemical activities, and macromolecular assemblies are coupled through mechanobiochemical feedback loops as depicted by the arrows in Fig. 1B.

**Figure 1.**
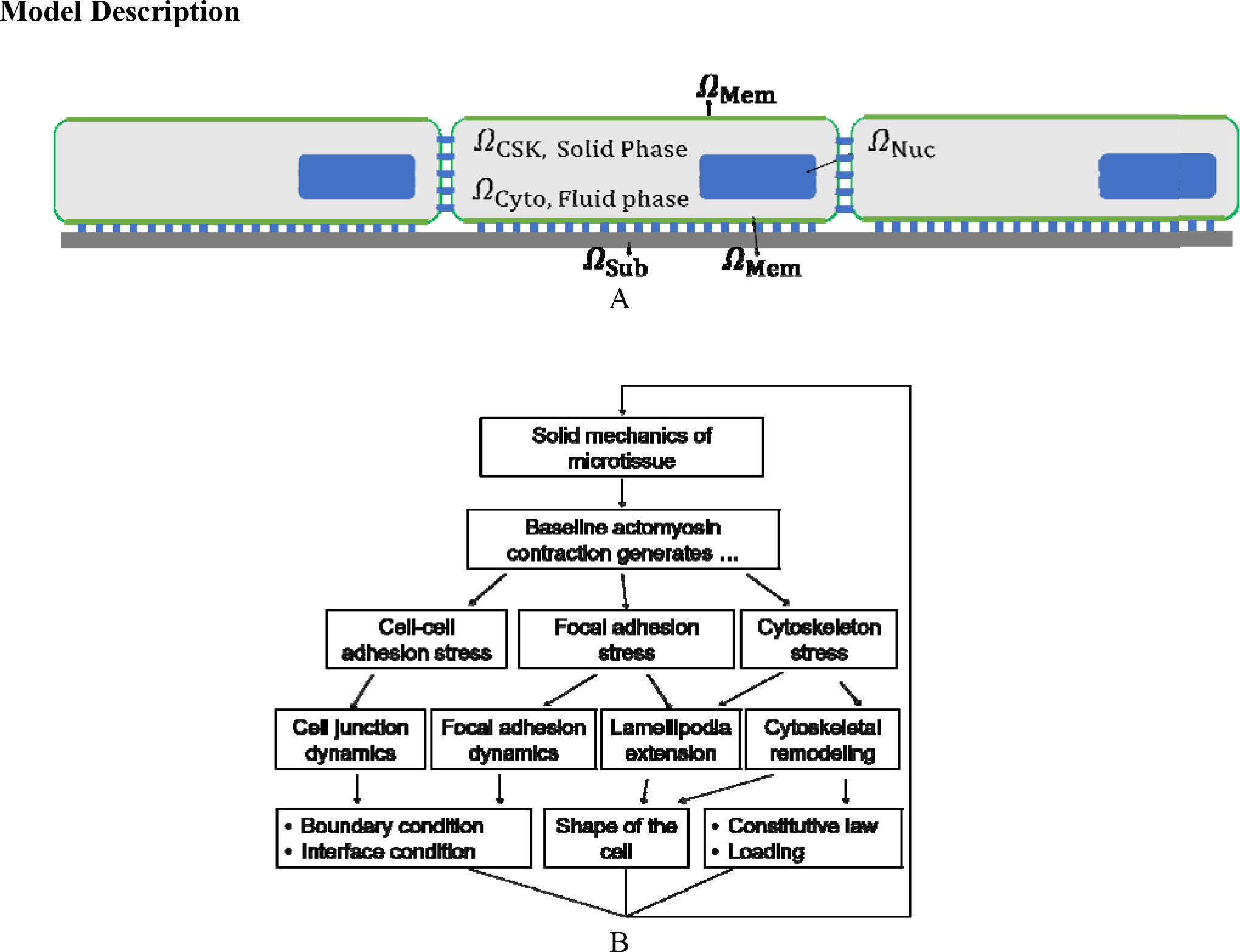
Schematic illustrations of (A) the physical domains in the cell model and Cell - Cell interaction (B) The mechanobiochemical coupling and feedback loops in the cell morphogenesis model.

#### Numerical Implementation of the model

The cell monolayer model is implemented in an in-house code using the finite element method, where Lagrangian mesh is adopted and 3-node triangle element is used. In the simulations of the movement of the cell, the nodal spatial coordinates are updated based on Eq. (10) and Eq. (11). An auto mesh-generating algorithm is utilized to perform re-meshing when mesh distortion occurs. Mesh transfer for the field variables is performed between the old and new mesh. Note that the parameter values used in the simulations presented below are chosen in a rather ad hoc fashion to give results in the same order of magnitude to experimental data in the literature. Use of parameter estimation algorithms to fit the parameters based on the known experimental data will be the subject of future studies.

### Simulation results

#### Establishment of polarity of the cell with the reaction-diffusion sub-model

Cell polarization (i.e., forming head and tail) is critical in cell migration to achieve directed movement. The spontaneous polarization has been thought as a pattern formation in reaction-diffusion systems ^31,51,57^ we here define the system of equations consisting of Eq. (6)-(9), where 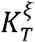, and 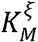 are set to be zero, as the reaction-diffusion sub-model. In this reaction-diffusion submodel, which were previously proposed by Maree et al ^32^, the protrusion signal *ζ* and retraction signal *ξ* inhibit each other through the 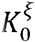 and 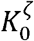 terms, respectively. As studied by Maree et al, this simple mutual-inhibition model can induce spontaneous polarization. Figure 2 shows the simulation result of the reaction-diffusion submodel. As shown in the top row of Fig 2, starting with a randomly perturbed initial state, the protrusion signal *ξ* (and the retraction signal, not shown) in a circular cell spontaneously polarizes, i.e., spatially separates into two zones: high and low regions.

**Figure 2.**
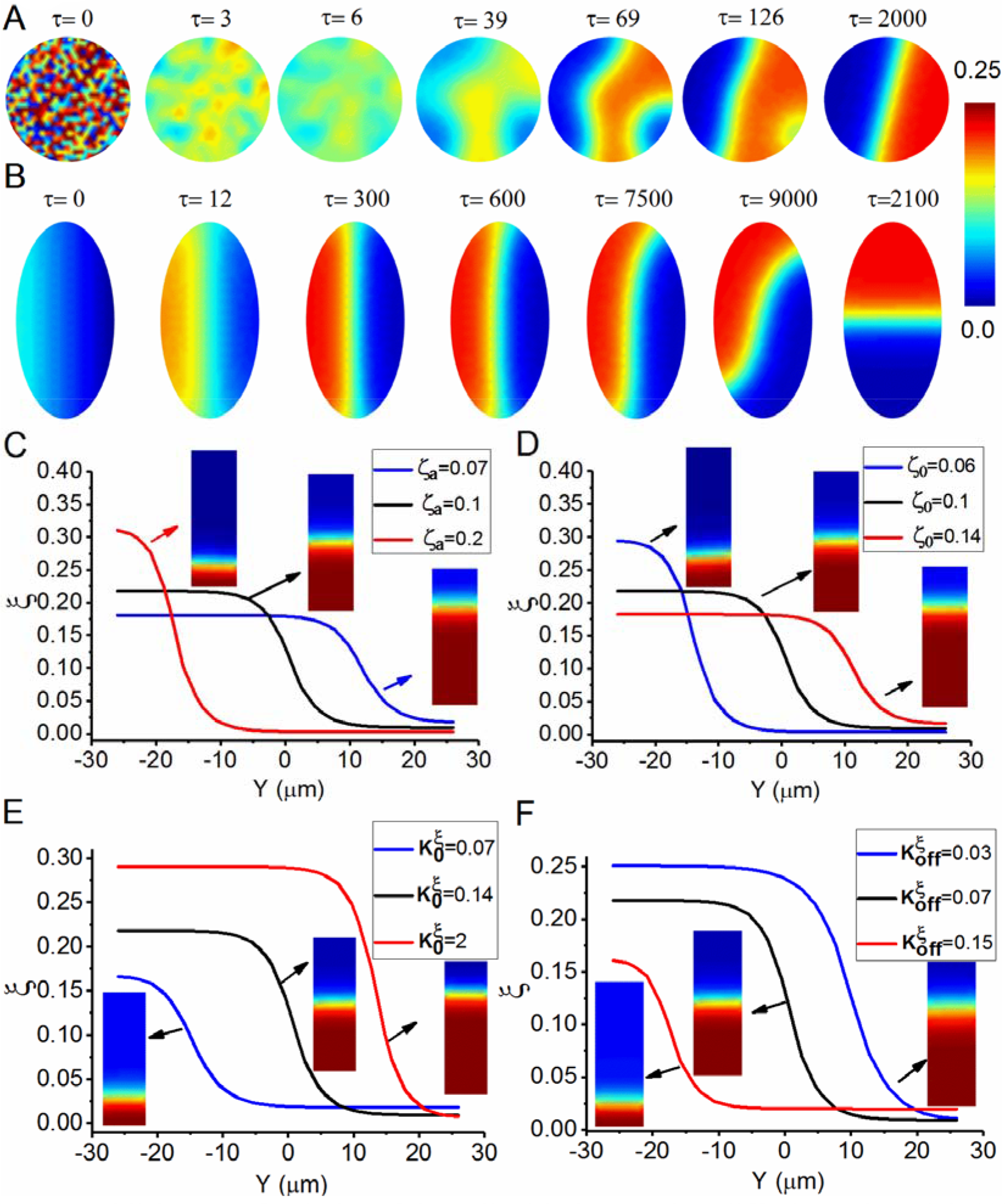
Establishment of polarity of the cell with the reaction-diffusion submodel. (A) In a circular cell, protrusion signal, *ξ*, polarizes spontaneously with random initial perturbation. (B) The effect of cell shape on the spatial patterns of protrusion signal in an elliptical cell. (C) Change in the protrusion signal distribution by varying the total concentration of the retraction signal, *ζ_a_*. (D) Change in the protrusion signal distribution by varying the model parameter, *ζ*_0_. (E) Change in the protrusion signal distribution by varying spontaneous activation parameter, 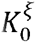. (F)Change in protrusion signal distribution by varying the decay parameter, 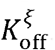. Parameter values used: 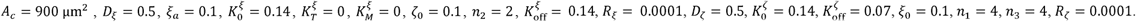 These parameter values are used for the remainder of the paper unless specifically mentioned.

As previously showed by Maree et al. ^51^, the cell shape (i.e., the shape of the mathematical domain of the reaction-diffusion equations) has an important effect on the spatial patterns formed. They concluded that at the steady state, the length of the interface that separates the high and low regions minimizes. Our simulation results agree with their conclusion. As shown in Fig. 2B, the interface in the elliptic cell is initially setup to be parallel to the longer axis of the ellipse (i.e., the initial distribution of the protrusion signal is a gradient from high in the left to low in the right). Overtime, the interface rotates and eventually aligns with the shorter axis of the ellipse. To further study the stable solutions of the reaction-diffusion submodel, a rectangular (with an aspect ratio of 1:3) shape is used to mimic approximately a quasi-1D version of this submodel. As shown in Fig. 2C-F, parameter *ζ*_*a*_, *ζ*_0_, 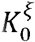, and 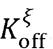 can shift the position of the interface and the peak value of *ξ* at the steady state. Note that the retraction signal distribution is the inverse of the of the protrusion (i.e., for the zone with high concentration of protrusion signals we observe low concentration of retraction signal and vice-versa). Fig 2C shows the effect of change in the protrusion signal distribution by varying the total retraction signal. This can be interpreted as increasing the total value of retraction signal will localize the distribution of the protrusion signal. In fig 2D we observe that increasing the model parameter, *ζ*_0_, amplifies the protrusion signal distribution. Fig 2E-2F shows the effect of variation of the spontaneous activation parameter, 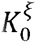 and the decay parameter, 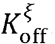, in protrusion signal distribution. It is intuitive that spontaneous activation parameter and decay parameter are inhibit each other, meaning the effect of increasing one of them is similar to the effect of the reducing of the other.

#### Focal adhesion stress-dependent protrusion signal distribution

The reaction-diffusion submodel described above can account for the polarization of the protrusion signal, but it cannot explain the phenomenon observed in the previous study ^25^ in which the membrane protrusion localized to the four corners of the square cell. Our previous studies showed that localization of the traction stress (which is equal to the focal adhesion stress) at the corners of the square cell is simply due to the mechanics principle of static equilibrium of an elastic body ^39^. Based on that, we argue that protrusion signal and focal adhesion assembly can be enhanced by the mechanical stress in the focal adhesion. We implement this hypothesis in the model by introducing the 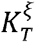 term in Eq. (6) and the 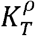 term in Eq. (16).

We define the system of equations consisting of Eq. (1)-(9) and Eq. (16)-(19), where 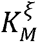. and 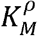 are set to be zero, as the stress-reaction-diffusion submodel. Simulations results of the stress-reaction-diffusion submodel for the square shape are presented in Fig. 3, showing the localization of high traction stress and protrusion signal at the corners of the square cell. As time goes on, the localization is enhanced by a positive feedback loop in the stress-reaction-diffusion submodel: larger traction stress *T* leads to bigger *ρ* (Eq. (16)), larger *ρ* leads to bigger *k_cs_* (Eq. (19)), bigger *k_cs_* results in larger traction stress in the continuum mechanics solution. Eventually, the traction stress distribution reaches to a steady state.

**Figure 3.**
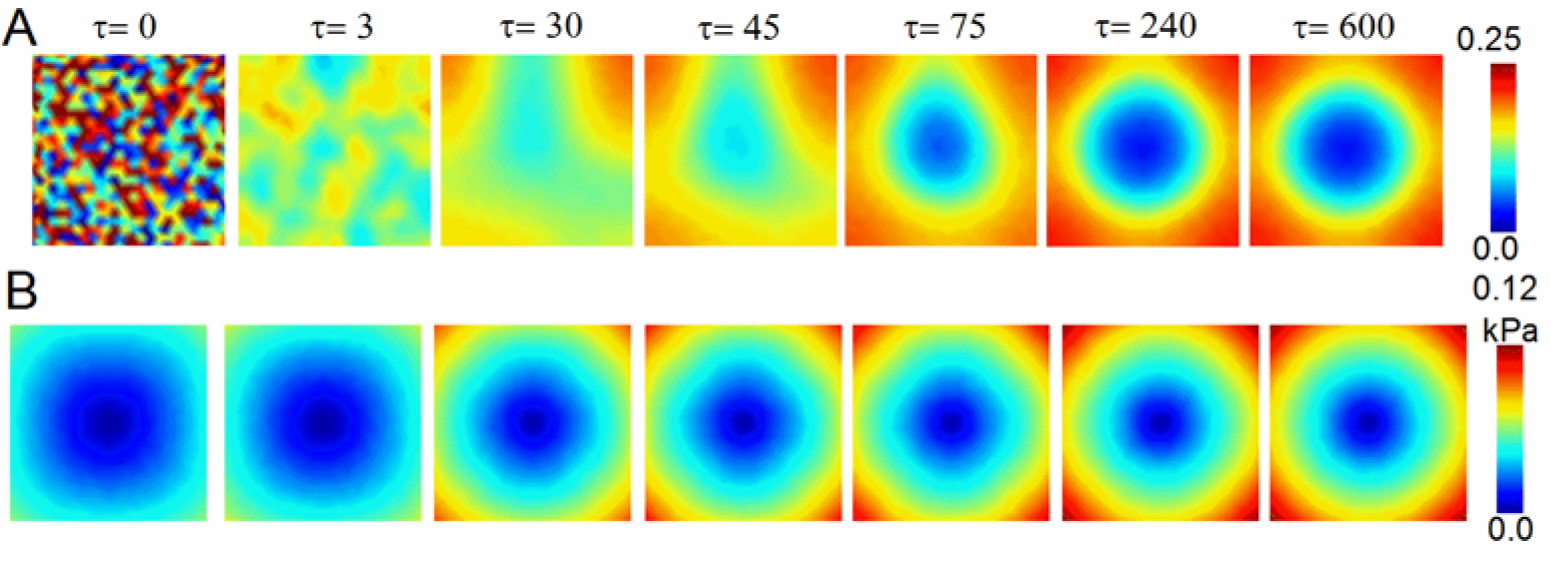
Coupling of the mechanical stresses of the cell to the protrusion signal (A) Protrusion signal, *ξ*, forms pattern spontaneously with random initial perturbation. (B) Traction stress contour, with coupling protrusion signal. (i.e., turning on stress dependent activation parameter, 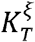). Parameter values used: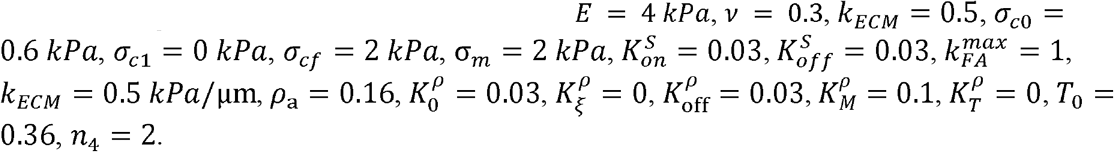.

#### The role of the cytoskeleton asymmetry to the persistent migration

The full model of the single cell, consisting of the cytoskeleton module, cell motility module, and the cell-substrate interaction module, describe the dynamic process of single-cell morphogenesis. Cell shape, which is one of the properties of cell morphology, is an emergent property of a cell spreading and migration. Cells of different types adopt different cell shapes during the spreading/migrating process.

In this model, we introduce a cytoskeleton-asymmetry function *f_M_*(*s*) to be able to explicitly control the directional persistence of migration. Figure 4 illustrates the role of function *f_M_*(*s*) in cell migration, in which the first two rows correspond to the model simulation where the cytoskeleton asymmetry is turned off (i.e., setting 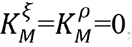), while the bottom two rows correspond to the simulation where the cytoskeleton asymmetry is on (see **Movie S1** and **S2**). Both simulations start with a circular cell and polarized protrusion signal. As shown in the first row, the cell first becomes an elliptic shape due to the protrusion on the front of the cell and the retraction on the back of the cell. The interface line that divides the high and low protrusion signal regions is parallel to the longer axis of the ellipse. Then the cell front turns due to the turning of the interface line of the protrusion signal towards the shorter axis of the ellipse as illustrated in Fig. 1B. Without the cytoskeleton asymmetry, the model cell moves but then turns. On the other hand, as shown in the third row of Fig. 4 where the cytoskeleton asymmetry is turned on, the cell shape becomes elongated in the dynamic equilibrium of the process of protrusion and retraction, and the cell preserves migration direction. With the cytoskeleton asymmetry, the model cell can preserve the migration direction.

**Figure 4.**
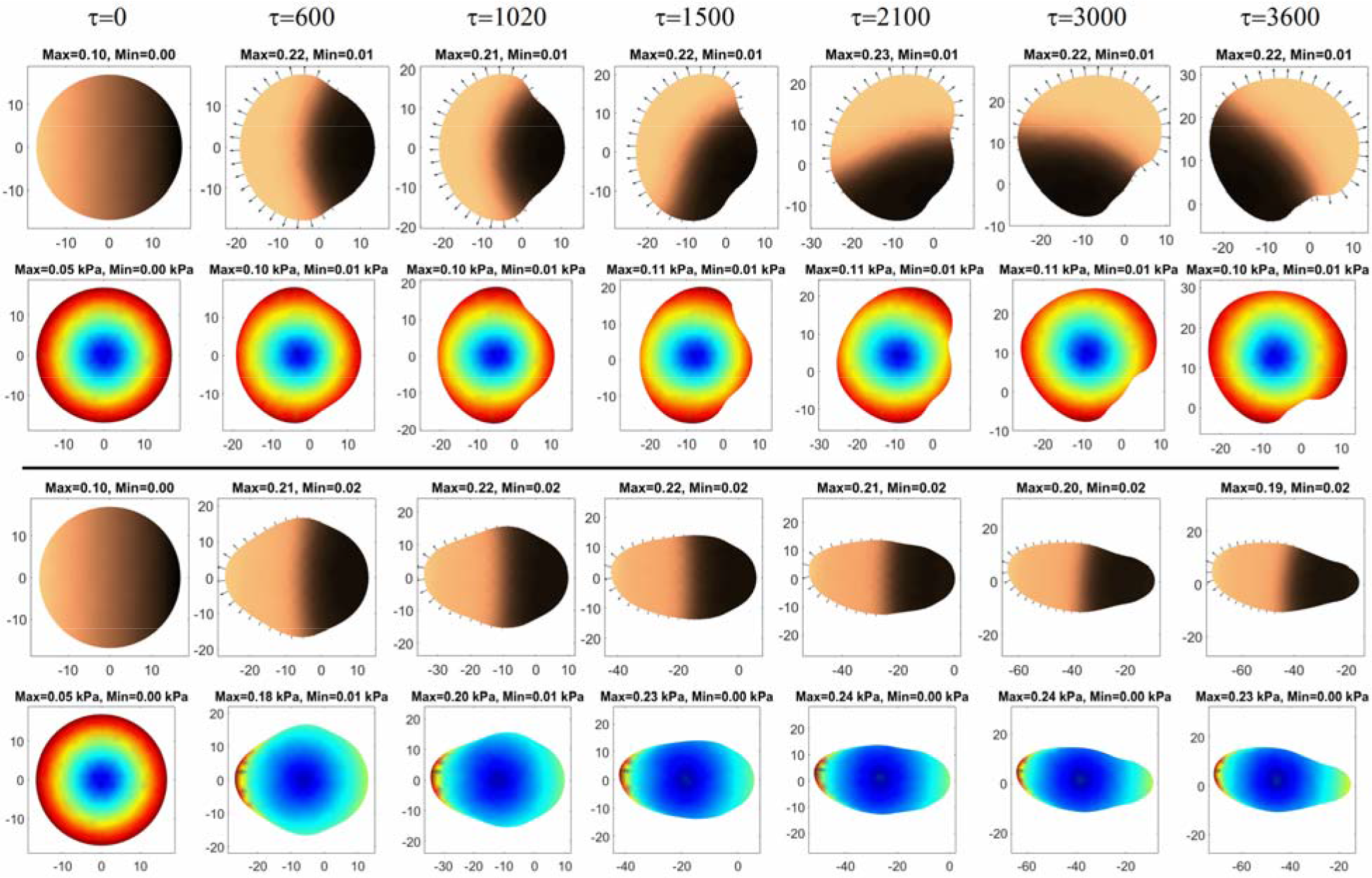
The role of the cytoskeleton asymmetry to the persistent migration. The first and third rows: protrusion signal. The second and fourth rows: traction stress. (also see **Movie S1** and **S2**). Parameter values used: for the first two rows: 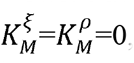, for the bottom two rows: 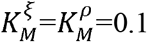. Following parameters are the same for both cases: *A*_max_ = 1800 μm^2^, *ξ*_*p*_ = 0, *c*_*p*_ = 0.25, *c_r_* = 0.01, *K_M_*= 0.02, *c_e_* = 10, *n* = 4.

#### Simulation of durotaxis

Durotaxis is a term coined by Lo et al. ^12^, which refers to the substrate rigidity-guided cell migration. They showed in the in vitro experiment where a fibroblast cell crawls from the stiffer side (i.e., the darker region) of the substrate toward the softer side (i.e., the brighter region), the cell made a 90-degree turn at the interface to avoid migrating into the softer region. A conceptual two-step theory consisting of the force generation and mechanotransduction has been proposed previously by Lo et al. to explain the durotaxis. A simple mechanics model has been presented by us previously to explain how exactly the larger focal adhesion stress is generated at the stiffer region of the substrate. Static equilibrium of the cell adhering to the substrate directly yield the disparate traction stress on regions of different rigidity^58^.

Simulations were setup to simulate durotaxis as a dynamic process. Cells started as a polarized circular shape and placed on the stiffer region of the substrate. The cell then crawls toward the softer region (i.e., left side). Snapshots from two simulations are presented in the first 2 rows of the Fig. 5 and bottom two rows of the Fig. 5. The only difference between these two simulations is the protrusion stress dependent activation parameter, 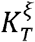, for the top two rows of Fig. 5,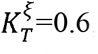, and for the bottom two rows, 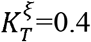. Parameter 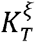 defines the level of focal adhesion stress-dependent activation of protrusion signal. At the larger value (Fig. 5 top two rows), the cell makes a turn at the stiff-to-soft interface, while at the smaller value, the cell crosses the interface (also see **Movie S3** and **S4**).

**Figure 5.**
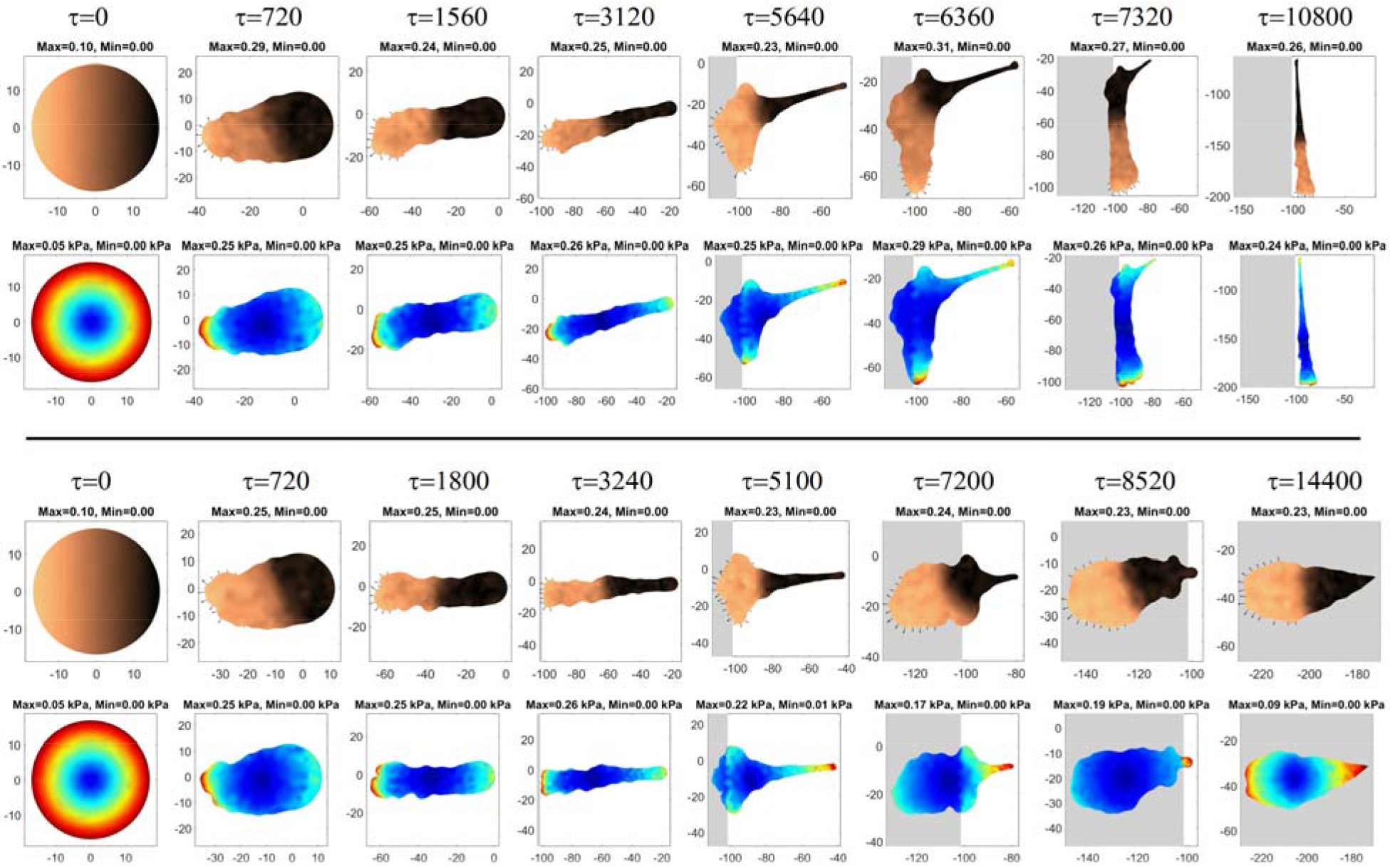
Simulation of durotaxis. (also see **Movie S3** and **S4**) The first and third rows: protrusion signal. The second and fourth rows: traction stress. Top two rows: higher stress dependent parameter, =0.6, durotaxis happens. Bottom two rows: lower stress dependent, =0.4, cell passes the interfaces.

#### Collective cell migration in monolayer tissue

Collective cell migration has been studied *in vitro* where a confluent monolayer of cells crawl on a flat 2d substrate ^59,60^. Here we conduct the *in-silico* modeling of cell crawling in monolayers, as shown in Fig. 6. Total of 26 cells are confined in an adhesive region of a circular shape and with a hole in the middle. The inner and outer radius of the adhesive region are 30 um and 83 um, respectively. To study the role of intercellular adhesion in collective cell migration, we performed two simulations: case-I (cell-cell adhesion is turned off) and case-II (cell-cell adhesion is on). The dynamic simulations start with circular cells seeded onto the adhesion region. Overtime, cells polarize, spread, and migrate (see **Movie S5** and **S6**).

**Figure 6.**
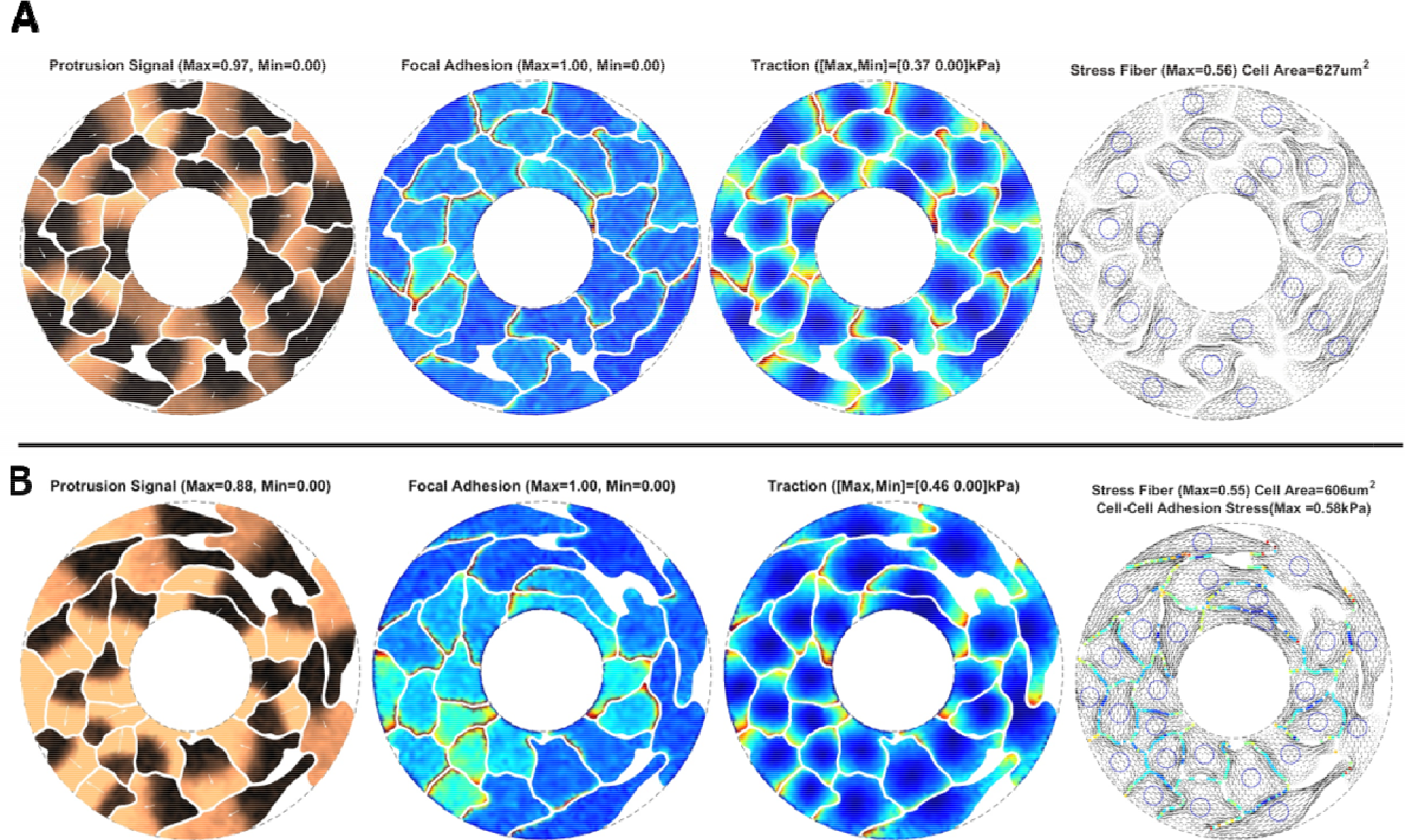
Collective cell migration in confluent monolayers. (A) Simulation snapshots of cells in confluent monolayers without cell-cell adhesion, (B) Simulation snapshots of cells in confluent monolayers with cell-cell adhesion. Four subfigures in each row show protrusion signal, focal adhesion, traction stress, and stress fiber, respectively (see movie S5, S6). In the stress fiber figures, fourth column, circles on the cells are the nucleus. Parameter value used:

Figure 6A and 6B show the simulation snapshots of cells in the confluent monolayers for case-I and case-II, respectively, where the migration direction of each cell can be seen in the
protrusion subfigures. The cell-cell adhesion stress is zero for case-I (Fig. 6A) since it is turned off. In case-II, because of the presence of the cell-cell adhesion, cell contraction is balanced by the cell-cell adhesion, rather than purely by the cell-substrate adhesion.

## Conclusions

In this work, we developed a 2D computational model for studying cell morphogenesis in monolayer tissues. Because of the complex nature of the living cell, the model, despite being phenomenological, is still sophisticated. Conceptually, we divide the full model into modules, and studied the behaviors of the submodels, as well as the couplings between modules. We have showed that the reaction-diffusion submodel can simulate cell polarization (head-to-tail formation), the stress-reaction-diffusion submodel can simulate the localization of protrusion signal to the corners of the square cell, and the cytoskeleton-asymmetry module can simulate the persistent migration. We also demonstrated that this mechanobiochemical model can simulate durotaxis and cell morphogenesis in monolayers.

Our computational model incorporates the reaction-diffusion equations with continuum mechanics equations, thus enabling in silico studies of the coupling between the biochemistry and mechanobiology. The finite element method-based numerical implementation of the model makes the computational model efficient in simulating cell monolayers with tens to hundreds of cells on desktop workstations. The computational model and the computer program can be used to test hypothesis and gain understandings of the complex system of living cells and tissues.

## Acknowledgments

H.Y. acknowledge funding support from the ASME Haythornthwaite Research Initiation Grant Award.

## References

1. Kang, H.-W. et al. A 3D bioprinting system to produce human-scale tissue constructs with structural integrity. Nat. Biotechnol. 34, 312–319 (2016).

2. Huh, D. et al. Reconstituting organ-level lung functions on a chip. Science 328, 1662–1668 (2010).

3. Murphy, S. V & Atala, A. 3D bioprinting of tissues and organs. Nat. Biotechnol. 32, 773–785 (2014).

4. Wyczalkowski, M. A., Chen, Z., Filas, B. A., Varner, V. D. & Taber, L. A. Computational Models for Mechanics of Morphogenesis. 152, 132–152 (2012).

5. Steinberg, M. S. Reconstruction of tissues by dissociated cells. Some morphogenetic tissue movements and the sorting out of embryonic cells may have a common explanation. Science 141, 401–8 (1963).

6. Marée, A. F. M., Hogeweg, P. & Mare, A. F. M. How amoeboids self-organize into a fruiting body: Multicellular coordination in Dictyostelium discoideum. Proc. Natl. Acad. Sci. 98, 3879–3883 (2001).

7. Computational Modeling in Tissue Engineering. (2013).

8. Davidson, E. H. et al. A genomic regulatory network for development. Science 295, 1669–1678 (2002).

9. Beloussov, L. V. The Dynamic Architecture of a Developing Organism: An Interdisciplinary Approach to the Development of Organisms. (1998).

10. Taber, L. A. Theoretical study of Beloussov’s hyper-restoration hypothesis for mechanical regulation of morphogenesis. Biomech. Model. Mechanobiol. 7, 427–441 (2008).

11. Howard, J., Grill, S. W. & Bois, J. S. Turing’s next steps: the mechanochemical basis of morphogenesis. Nat Rev Mol Cell Biol 12, 392–398 (2011).

12. Lo, C. M., Wang, H. B., Dembo, M. & Wang, Y.-L. Cell movement is guided by the rigidity of the substrate. Biophys. J. 79, 144–152 (2000).

13. Uttayarat, P., Toworfe, G. K., Dietrich, F., Lelkes, P. I. & Composto, R. J. Topographic guidance of endothelial cells on silicone surfaces with micro-to nanogrooves: Orientation of actin filaments and focal adhesions. J. Biomed. Mater. Res. Part A 75A, 668–680 (2005).

14. Den Braber, E. T., De Ruijter, J. E., Ginsel, L. A., Von Recum, A. F. & Jansen, J. A. Quantitative analysis of fibroblast morphology on microgrooved surfaces with various groove and ridge dimensions. Biomaterials 17, 2037–2044 (1996).

15. Oakley, C., Jaeger, N. A. F. & Brunette, D. M. Sensitivity of fibroblasts and their cytoskeletons to substratum topographies: Topographic guidance and topographic compensation by micromachined grooves of different dimensions. Exp. Cell Res. 234, 413–424 (1997).

16. van Delft, F. et al. Manufacturing substrate nano-grooves for studying cell alignment and adhesion. in 85, 1362–1366 (Elsevier Science Bv, 2008).

17. denBraber, E. T., deRuijter, J. E., Ginsel, L. A., vonRecum, A. F. & Jansen, J. A. Quantitative analysis of fibroblast morphology on microgrooved surfaces with various groove and ridge dimensions. Biomaterials 17, 2037–2044 (1996).

18. Clark, P. et al. Topographical Control of Cell Behavior .2. Multiple Grooved Substrata. Development 108, 635–644 (1990).

19. Harunaga, J. S. & Yamada, K. M. Cell-matrix adhesions in 3D. Matrix Biol. 30, 363–368 (2011).

20. Grinnell, F., Ho, C.-H., Tamariz, E., Lee, D. J. & Skuta, G. Dendritic Fibroblasts in Threedimensional Collagen Matrices. Mol. Biol. Cell 14, 384–395 (2003).

21. Grinnell, F. Fibroblast biology in three-dimensional collagen matrices. Trends Cell Biol. 13, 264–269 (2003).

22. Baker, B. M. & Chen, C. S. Deconstructing the third dimension - how 3D culture microenvironments alter cellular cues. J. Cell Sci. 125, 3015–3024 (2012).

23. Doyle, A. D., Wang, F. W., Matsumoto, K. & Yamada, K. M. One-dimensional topography underlies three-dimensional fi brillar cell migration. J. Cell Biol. 184, 481–490 (2009).

24. Chen, C. S., Tan, J. & Tien, J. MECHANOTRANSDUCTION AT CELL-MATRIX AND CELL-CELL CONTACTS. Annu. Rev. Biomed. Eng. 6, 275–302 (2004).

25. Parker, K. K. et al. Directional control of lamellipodia extension by constraining cell shape and orienting cell tractional forces. Faseb J 16, 1195–1204 (2002).

26. Grosberg, A. et al. Self-Organization of Muscle Cell Structure and Function. PLoS Comput Biol 7, e1001088 (2011).

27. Parker, K. K., Tan, J., Chen, C. S. & Tung, L. Myofibrillar architecture in engineered cardiac myocytes. Circ Res 103, 340–342 (2008).

28. Bray, M. A., Sheehy, S. P. & Parker, K. K. Sarcomere alignment is regulated by myocyte shape. Cell Motil. Cytoskeleton 65, 641–651 (2008).

29. Onsum, M. D. & Rao, C. V. Calling heads from tailsↃ: the role of mathematical modeling in understanding cell polarization. 74–81 (2009). doi:10.1016/j.ceb.2009.01.001

30. Wedlich-Soldner, R., Altschuler, S., Wu, L. & Li, R. Spontaneous cell polarization through actomyosin-based delivery of the Cdc42 GTPase. Science (80-.). 299, 1231–1235 (2003).

31. Satulovsky, J., Lui, R. & Wang, Y. Exploring the Control Circuit of Cell Migration by Mathematical Modeling. Biophys. J. 94, 3671–3683 (2008).

32. Marée, A. F. M., Jilkine, A., Dawes, A., Grieneisen, V. A. & Edelstein-Keshet, L. Polarization and movement of keratocytes: A multiscale modelling approach. Bulletin of Mathematical Biology 68, (2006).

33. Novak, I. L., Slepchenko, B. M., Mogilner, A. & Loew, L. M. Cooperativity between Cell Contractility and Adhesion. Phys. Rev. Lett. 93, 268109 (2004).

34. Zeng, X. & Li, S. Multiscale modeling and simulation of soft adhesion and contact of stem cells. J. Mech. Behav. Biomed. Mater. 4, 180–189 (2011).

35. Deshpande, V. S., McMeeking, R. M. & Evans, A. G. A bio-chemo-mechanical model for cell contractility. Proc. Natl. Acad. Sci. U. S. A. 103, 14015–14020 (2006).

36. Walcott, S. & Sun, S. X. A mechanical model of actin stress fiber formation and substrate elasticity sensing in adherent cells. Proc. Natl. Acad. Sci. U. S. A. 107, 7757–7762 (2010).

37. Kang, J. et al. Response of an actin filament network model under cyclic stretching through a coarse grained Monte Carl approach. J. Theor. Biol. 274, 109–119 (2011).

38. Pathak, A., Deshpande, V. S., McMeeking, R. M. & Evans, A. G. The simulation of stress fibre and focal adhesion development in cells on patterned substrates. J. R. Soc. Interface 5, 507–524 (2008).

39. Yuan, H., Marzban, B. & Parker, K. K. Myofibrils in Cardiomyocytes Tend to Assemble Along the Maximal Principle Stress Directions. J Biomech Eng doi: 10.11, (2017).

40. Forgacs, G. & Newman, S. A. Biological Physics of the Developing Embryo. (Cambridge University Press, 2005). doi:DOI: 10.1017/CBO9780511755576

41. Ridley, A. J. et al. Cell migration: integrating signals from front to back. Science 302, 1704–9 (2003).

42. Polacheck, W. J., Zervantonakis, I. K. & Kamm, R. D. Tumor cell migration in complex microenvironments. Cell. Mol. Life Sci. 13, 1133–1145 (2012).

43. Friedl, P., Sahai, E., Weiss, S. & Yamada, K. M. New dimensions in cell migration. Nat. Rev. Mol. Cell Biol. 13, 743 (2012).

44. Devreotes, P. & Horwitz, A. R. Signaling networks that regulate cell migration. Cold Spring Harb Perspect Biol 7, a005959 (2015).

45. Danuser, G., Allard, J. & Mogilner, A. Mathematical modeling of eukaryotic cell migration: insights beyond experiments. Annu. Rev. Cell Dev. Biol. 29, 501–28 (2013).

46. Iglesias, P. a. & Devreotes, P. N. Biased excitable networks: How cells direct motion in response to gradients. Curr. Opin. Cell Biol. 24, 245–253 (2012).

47. Xiong, Y., Huang, C.-H., Iglesias, P. A. & Devreotes, P. N. Cells navigate with a local-excitation, global-inhibition-biased excitable network. Proc. Natl. Acad. Sci. 107, 17079–17086 (2010).

48. Li, S. et al. The role of the dynamics of focal adhesion kinase in the mechanotaxis of endothelial cells. Proc. Natl. Acad. Sci. U. S. A. 99, 3546–3551 (2002).

49. Polacheck, W. J., German, A. E., Mammoto, A., Ingber, D. E. & Kamm, R. D. Mechanotransduction of fluid stresses governs 3D cell migration. Proc. Natl. Acad. Sci. U. S. A. 111, 2447–52 (2014).

50. Trepat, X. & Fredberg, J. J. Plithotaxis and emergent dynamics in collective cellular migration. Trends Cell Biol. 21, 638–646 (2011).

51. Marée, A. F. M., Grieneisen, V. A. & Edelstein-Keshet, L. How cells integrate complex stimuli: The effect of feedback from phosphoinositides and cell shape on cell polarization and motility. PLoS Comput. Biol. 8, (2012).

52. Prentice-Mott, H. V. et al. Directional memory arises from long-lived cytoskeletal asymmetries in polarized chemotactic cells. Proc. Natl. Acad. Sci. 113, 201513289 (2016).

53. McCain, M. L., Lee, H., Aratyn-Schaus, Y., Kléber, A. G. & Parker, K. K. Cooperative coupling of cell-matrix and cell–cell adhesions in cardiac muscle. Proc. Natl. Acad. Sci. 109, 9881–9886 (2012).

54. Aratyn-Schaus, Y. et al. Coupling primary and stem cell–derived cardiomyocytes in an in vitro model of cardiac cell therapy. J. Cell Biol. February 8, (2016).

55. Chopra, A., Tabdanov, E., Patel, H., Janmey, P. a & Kresh, J. Y. Cardiac myocyte remodeling mediated by N-cadherin-dependent mechanosensing. Am. J. Physiol. - Hear. Circ. Physiol. 300, H1252–H1266 (2011).

56. Sim, J. Y. et al. Spatial distribution of cell-cell and cell-ECM adhesions regulates force balance while main taining E-cadherin molecular tension in cell pairs. Mol. Biol. Cell 26, 2456–2465 (2015).

57. Altschuler, S. J., Angenent, S. B., Wang, Y. & Wu, L. F. On the spontaneous emergence of cell polarity. Nature 454, 886–889 (2008).

58. Marzban, B., Yi, X. & Yuan, H. A minimal mechanics model for mechanosensing of substrate rigidity gradient in durotaxis. Biomech. Model. Mechanobiol. (2018). doi:10.1007/s10237-018-1001-3

59. Rolli, C. G. et al. Switchable adhesive substrates: Revealing geometry dependence in collective cell behavior. Biomaterials 33, 2409–2418 (2012).

60. Tambe, D. T. et al. Collective cell guidance by cooperative intercellular forces. Nat. Mater. 10, 469–75 (2011).

